# Reactivities of the Front Pocket N-Cap Cysteines in Human Kinases

**DOI:** 10.1101/2021.06.28.450170

**Authors:** Ruibin Liu, Shaoqi Zhan, Ye Che, Jana Shen

## Abstract

Discovery of targeted covalent inhibitors directed at nucleophilic cysteines is attracting enormous interest. The front pocket (FP) N-cap cysteine has been the most popular site of covalent modification in kinases. Curiously, a long-standing hypothesis associates the N-cap position with cysteine hyper-reactivity; however, traditional computational methods suggest that the FP N-cap cysteines in all human kinases are predominantly unreactive at physiological pH. Here we applied a newly developed GPU-accelerated continuous constant pH molecular dynamics (CpHMD) tool to test the N-cap hypothesis and elucidate the cysteine reactivities. Simulations showed that the N-cap cysteines in BTK/BMX/TEC/ITK/TXK, JAK3, and MKK7 sample the reactive thiolate form to varying degrees at physiological pH; however, those in BLK and EGFR/ERBB2/ERBB4 which contain an Asp at the N-cap+3 position adopt the unreactive thiol form. The latter argues in favor of the base-assisted thiol-Michael addition mechanisms as suggested by the quantum mechanical calculations and experimental structure-function studies of EGFR inhibitors. Analysis revealed that the reactive N-cap cysteines are stabilized by hydrogen bond as well as electrostatic interactions, and in their absence a N-cap cysteine is unreactive due to desolvation. To test a corollary of the N-cap hypothesis, we also examined the reactivities of the FP N-cap+2 cysteines in JNK1/JNK2/JNK3 and CASK. Additionally, our simulations predicted the reactive cysteine and lysine locations in all 15 kinases. Our findings offer a systematic understanding of cysteine reactivities in kinases and demonstrate the predictive power and physical insights CpHMD can provide to guide the rational design of targeted covalent inhibitors.

## INTRODUCTION

Protein kinases are enzymes that catalyze protein phosphorylation reactions, and represent one of the largest gene families with over 500 genes discovered so far. ^1^ Dysregulation of kinase functions plays significant roles in cancer, immunology, and a host of other diseases. ^1^ In the past decade, the highest number of approvals by the US Food and Drug Administration (FDA) have gone to inhibitors targeting kinases as compared to other protein families. ^2^ While traditional kinase inhibitors are small molecules that bind the ATP-binding site via reversible interactions, ^1^ targeted covalent kinase inhibitors (TCKIs) are emerging at a rapid pace ^3,4^ due to the advantages such as prolonged duration of action, enhanced potency, and increased selectivity. ^5,6^ Starting from a low affinity reversible inhibitor, a TCKI is typically developed by structure-guided incorporation of an electrophilic warhead that covalently links with a nearby nucleophilic side chain. ^5^

Owing to the high nucleophilicity of thiolate and the non-catalytic role in kinases, cysteine is the most targeted amino acid in covalent drug discoveries. ^1,5,21^ According to a survey from 2018, ^3^ over half of the TCKIs are directed at a Cys located in the linker at the N-terminal end (N-cap) of the *α*D helix near the front pocket (FP, Fig. 1). Sequence alignment (by us and others ^5,22^) showed that 11 kinases in the human kinome possess such a FP N-cap Cys, including BTK/BMX/TEC/ITK/TXK from the Tec family, BLK from the Src family, JAK3 from the JakA family, EGFR/ERBB2/ERBB4 from the EGFR family, and MKK7 from the STE7 family (Fig. 1), all of which are therapeutic targets.

**Figure 1.**
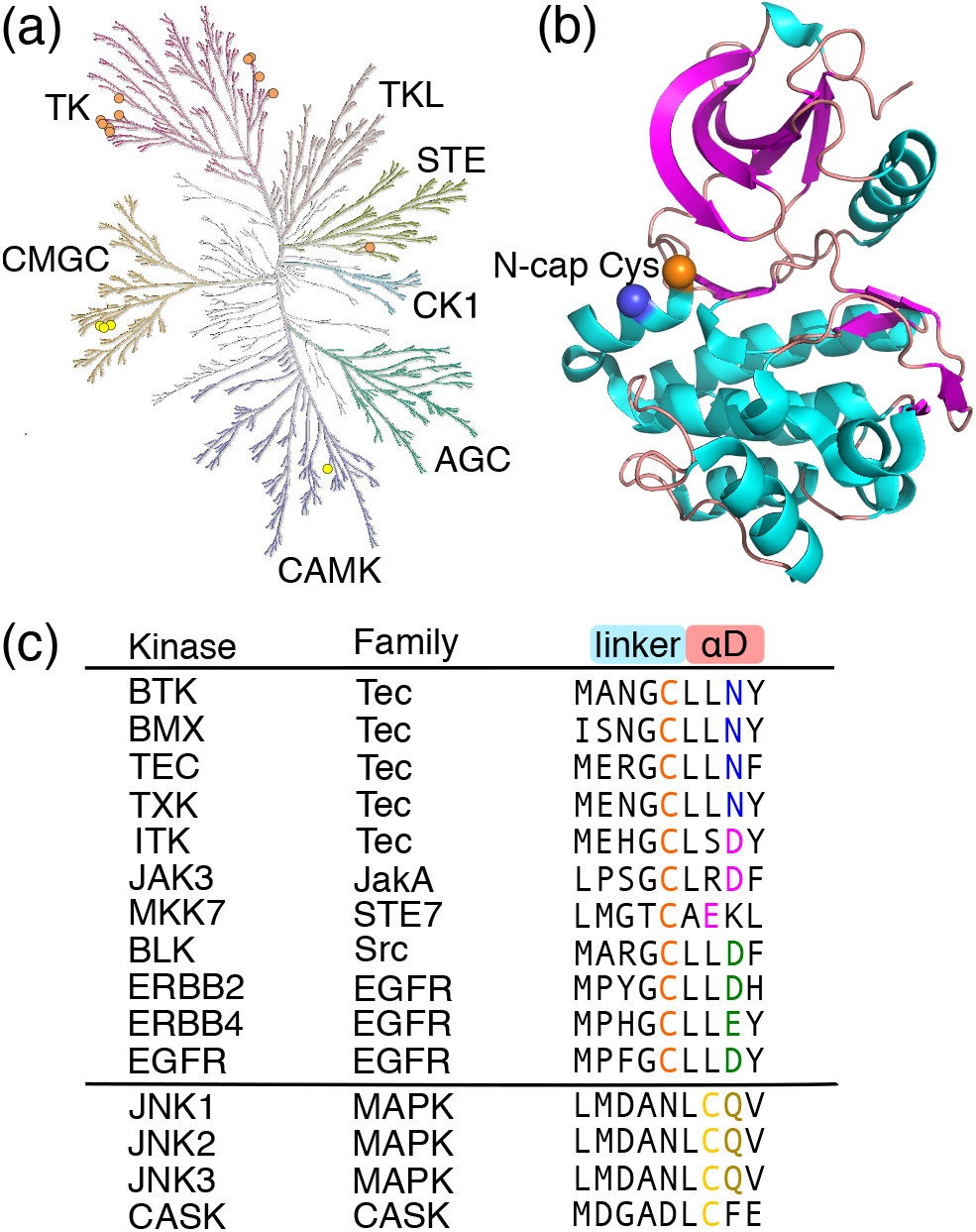
Human kinases with a FP N-cap or N-cap+2 Cys. (a) 15 kinases possessing a FP N-cap or N-cap+2 Cys displayed in the human kinome tree (generated with KinMap ^7^). (b) Structure of the kinase domain of BMX showing a FP N-cap Cys (orange sphere, PDB ID: 3sxs ^8^). The frequently occurring Asn/Asp/Glu at N-cap+3 position is shown as a blue sphere. (c) Names of the kinases possessing a FP N-cap or N-cap+2 Cys, the families they belong to, and the sequences of nearby amino acids. The FP N-cap Cys is colored orange, the N-cap+2 Cys is colored yellow, and nearby residues making a significant contribution to its p*K* _a_ shift are colored blue, magenta, green, or brown (same as in Fig. 2).

BTK/BMX/TEC/ITK/TXK are key components of T-cell receptor signaling and contribute to acting reorganization and cell polarization. ^8,23–25^ BLK is an oncogene and a potential therapy target in cutaneous T-cell lymphoma. ^26^ JAK3 drives signaling through cytokine receptors, and is associated with various immune-mediated diseases. ^27,28^ EGFR (or ERBB1 or HER1), ERBB2 (or HER2), and ERBB4 (or HER4) play essential roles in regulating cell proliferation, survival, differentiation and migration, and understanding their roles has fueled the development of targeted therapies for cancer in the past decade. ^29–32^ MKK7 is associated with multiple myeloma and various neurological diseases. ^15,16,33^

The FP N-cap Cys in all human kinases have been covalently modified. Most notably, the FDA-approved drug ibrutinib covalently modifies the N-cap Cys in BTK/BMX/TEC/ITK, BLK, JAK3, and EGFR/ERBB4 (Table 1). ^22,34,35^ The same Cys in EGFR is also targeted by newer covalent drugs such as afatinib, neratinib, acalabrutinib, spebrutinib, mavelertinib, and osimertinib. The popularity of the FP N-cap Cys in TCKI discovery can be explained by its accessibility (near the ATP binding site) and expected reactivity (see below). A N-cap Asp has been known to stabilize a helix, which was attributed to the favorable interaction between the negative aspartate and the positive helix dipole. ^36,37^ Similarly, a N-cap Cys was found to be helix stabilizing, which was attributed to the thiolate-helix dipole interaction. ^38,39^ It was argued that Cys is the least frequent N-cap residue, because it is in the thiolate form prone to chemical modification. ^38^ Indeed, the N-cap Cys in myoglobin has a measured p*K* _a_ of 6.5, which is 1.9 or 1.6 units lower than the (mutant) Cys at N2 or N1 position, and 2 units lower than the p*K* _a_ of 8.6 for the model Cys in a peptide. ^39^ Thus, the consensus is that a N-cap Cys is in the hyper-reactive thiolate form at physiological pH, which we will refer to as the N-cap hypothesis.

**Table 1.** Summary of kinases with a FP N-cap or N-cap+2 Cys and the covalent inhibitors*^a^* directed at the location

| Kinase | Family | Group | PDB | Residue | PROPKA<sup>b</sup> | TCKI |
| --- | --- | --- | --- | --- | --- | --- |
| **FP N-cap Cys** |  |  |  |  |  |  |
| BTK | Tec | TK | 3pj3 | C481 | 9.6 | ibrutinib, acalabrutinib, zanubrutinib, tirabrutinib,<sup>9</sup> orelabrutinib<sup>10</sup> |
| BMX | Tec | TK | 3sxs | C496 | 9.9 | ibrutinib |
| TEC | Tec | TK | 4hct<sup>a</sup> | C449 | 9.2 | ibrutinib, ritlecitinib<sup>c</sup> |
| ITK | Tec | TK | 3miy | C442 | 10.7 | ibrutinib |
| TXK | Tec | TK | 3t9t<sup>a</sup> | C350 | 10.4 | acalabrutinib |
| BLK | Src | TK | 4mxy<sup>a</sup> | C319 | 10.6 | ibrutinib, acalabrutinib |
| JAK3 | JakA | TK | 5lwm | C909 | 10.3 | ritlecitinib<sup>11</sup> |
| EGFR | EGFR | TK | 4zjv | C797 | 11.4 | afatinib, osimertinib, dacomitinib, neratinib, olmutinib,<sup>12</sup> pyrotinib,<sup>13</sup> mobocertinib,<sup>14</sup> ibrutinib, acalabrutinib |
| ERBB2 | EGFR | TK | 3pp0 | C805 | 10.8 | acalabrutinib |
| ERBB4 | EGFR | TK | 2r4b | C803 | 11.0 | ibrutinib |
| MKK7 | STE7 | STE | 6qfr | C218 | 9.1 | investigational inhibitors<sup>15,16</sup> |
| **FP N-cap+2 Cys** |  |  |  |  |  |  |
| JNK1 | MAPK | CMGC | 2xrw | C116 | 10.0 | investigational inhibitors<sup>17</sup> |
| JNK2 | MAPK | CMGC | 3e7o | C116 | 10.2 | investigational inhibitors<sup>17</sup> |
| JNK3 | MAPK | CMGC | 6emh | C154 | 10.4 | investigational inhibitors<sup>18</sup> |
| CASK | CASK | CAMK | 3mfr | C100 | 13.0 | unknown |

<sup>a</sup> Inhibitors without references are from the FDA approved ones from the FDA Orange Book (https://www.accessdata.fda.gov/scripts/cder/ob/index.cfm). Those with references are in the last stage of clinical trial; some of them have gained regulatory approval in foreign countries. <sup>b</sup> PROPKA calculations were performed with PROPKA3.<sup>19</sup>
<sup>c</sup> Template structure used for homology modeling with SWISS-MODEL.<sup>20</sup>

Kinetics experiments demonstrated that the observed (apparent) rate constants of thiol reactions with electrophiles are inversely related to the thiol p*K* _a_’s. ^40–43^ Thus, it is generally accepted that cysteines with low p*K* _a_’s are more prone to covalent modification due to the increased availability of the nucleophilic thiolate form, and the Cys p*K* _a_ can be used as a proxy for reactivity in TCKI design. ^1,5,21,38^ Curiously, however, according to the popular empirical p*K* _a_ prediction program PROPKA, ^19^ the N-cap Cys in the aforementioned 11 kinases have p*K* _a_’s of 9.1–11.4 (Table 1), suggesting that they are mainly in the thiol form and consequently unreactive or have low reactivities towards electrophiles. Note, the intrinsic nucleophilicity of a thiolate often increases and sometimes stays constant or decreases with the thiol p*K* _a_ value depending on the molecule by molecule basis; ^40,43^ this is not a topic of this paper.

Accurate prediction of Cys p*K* _a_’s is challenging using traditional computational methods. For a dataset of 18 proteins, Rowley and coworker showed that the root-mean-square errors (RMSE’s) from the experimental p*K* _a_’s are 3.4–4.7 using the static structure based Poisson-Boltzmann (PB) and empirical PROPKA calculations. ^44^ They obtained a RMSE of 2.4–3.2 using the molecular dynamics (MD) based thermodynamic integration (TI) method, which is several orders of magnitude slower than the PB or PROKA calculations. This level of accuracy is however on par with that (RMSE of 2.7) from the null model which assumes the solution (or model) p*K* _a_ of 8.6^45^ for all cysteines.

Recently, we developed a GPU-accelerated continuous constant pH MD (CpHMD) method in Amber package ^46^ for accurate and rapid prediction of protein p*K* _a_’s based on independent pH ^47^ or pH replica-exchange titration ^48^ simulations in the GB-Neck2 implicit solvent. ^49^ Using replica-exchange CpHMD, which significantly accelerates the p*K* _a_ convergence, we scanned the kinome structure database and found that the catalytic (roof) Lys in a dozen of human kinases can become reactive in the rare DFG-out/*α*C-out inactive conformation. ^50^ Most recently, we implemented an asynchronous algorithm to allow replica-exchange simulations on a single or any number of GPU cards. ^51^ Benchmark study based on 24 proteins, including those with large p*K* _a_ downshifts examined by Rowley and coworker ^44^ and additional ones with large p*K* _a_ upshifts relative to the model value, gave a RMSE of 1.2, which is more than two units lower than the PB or empirical calculations. The accuracy of predicting thiolates at physiological pH is about 81% with the replica-exchange CpHMD, in contrast to the accuracy below 50% with the PB or empirical methods. ^52^

Motivated by the importance of the N-cap Cys in TCKI design and intrigued by the contradiction between the N-cap hypothesis and the high p*K* _a_’s predicted by the empirical method, here we employed the asynchronous replica-exchange CpHMD to determine the p*K* _a_’s of all 11 human kinases that possess a FP N-cap Cys. We found that most N-cap Cys have p*K* _a_’s in the range of 7.7–8.5, thus providing nucleophilic thiolates for the direct thiol-Michael addition with electrophiles at physiological pH; however, surprisingly, the N-cap Cys in EGFR/ERBB2/ERBB4 and BLK remain in the thiol form up to pH 10.5. Analysis of the pH-dependent conformational environment surrounding the N-cap Cys provides insights into the varied reactivities and support to the general base assisted mechanism for the Michael addition with the N-cap Cys in EGFR/ERBB2/ERBB4 and BLK. To further explore the N-cap hypothesis, we also determined the p*K* _a_’s of the N-cap+2 Cys in JNK1/JNK2/JNK3 and CASK. Additionally, our study identified reactive Cys and Lys locations in all 15 kinases, offering new opportunities for TCKI discovery. Together, our findings offer a systematic understanding of the Cys structure-reactivity relationship and demonstrate the utility of CpHMD simulations in aiding the rational design of Cys- and Lys-targeted covalent kinase inhibitors.

## RESULTS AND DISCUSSION

### CpHMD titration showed varied reactivities for the FP N-cap Cys

We performed the GPU-accelerated replica-exchange CpHMD titration simulations ^47,48,51^ to determine the Cys and Lys p*K* _a_ values for the aforementioned 11 human kinases possessing a FP N-cap Cys, which is located in the linker at the N-terminal end of the *α*D helix (position 52 of the ATP binding site sequence according to KLIFS database ^53^). Except for TEC, TXK, and BLK, for which homology models were built using SWISS-MODEL, ^20^ simulations started from the X-ray crystal structures with inhibitors, ions, and crystal water molecules removed. A pH range 6.0-10.5 was used, and each replica lasted 30–50 ns until convergence (for more details see Methods). Considering the inverse relationship between the thiol reactivity and p*K* _a_ ^40,54^ and to facilitate discussion, we group the Cys (or later Lys) in four categories (Table 2): unreactive (p*K* _a_ > 9.4), somewhat reactive (p*K* _a_ 8.4–9.4), reactive (p*K* _a_ 7.4–8.4), and hyper-reactive p*K* _a_ < 7.4. Accordingly, the fraction of thiolate (or neutral lysine) state at physiological pH is < 1% (unreactive), 1–10% (somewhat reactive), 10–50% (reactive), and > 50% (hyper-reactive).

**Table 2.** Reactivity definitions used in this work

| Cys/Lys $pK_a$ | Deprot. fraction | Reactivity |
| --- | --- | --- |
| $< 7.4$ | $> 50\%$ | hyper reactive |
| 7.4–8.4 | 10–50% | reactive |
| 8.5–9.4 | 1–10% | somewhat reactive |
| $> 9.4$ | $< 1\%$ | unreactive |

Our simulations gave a wide p*K* _a_ range from 7.6 to above 10.5 for the FP N-cap Cys, suggesting that its reactivity varies despite the same N-cap location. Among all, the N-cap Cys in BTK/BMX of the Tec family and JAK3 of the JakA family are nearly hyper-reactive, with the calculated p*K* _a_’s around 7.7 (Fig. 2 and Table S1). The N-cap Cys in TEC/TXK/ITK of the Tec family are also reactive, with the calculated p*K* _a_’s around 8.4 (Fig. 2 and Table S1), close to the value (8.6) for the model alanine penta-peptide (AACAA) in solution. ^45^ The N-cap Cys in MKK7 is somewhat reactive with a calculated p*K* _a_ of 9.1 (Fig. 2 and Table S1). Interestingly, the N-cap Cys in EGFR/ERBB2/ERBB4 of the EGFR family and BLK of the Src family are predicted as unreactive with p*K* _a_ values near or above 10.5 (Fig. 2 and Table S1), in agreement with our previous work. ^50^ Noticeably, our calculated p*K* _a_’s of 7.7 and 8.4 for the FP N-cap Cys in BTK and ITK are in quantitative agreement with the experimental values of 7.7 and 8.5, respectively. ^55^ However, curiously, our simulation showed that EGFR’s N-cap Cys remained protonated in the pH range 7.5– 10.5, in disagreement with the p*K* _a_ of 5.5 estimated based on the Bromobimane fluorescence labeling experiment. ^56^ We will come back to the discussion.

**Figure 2.**
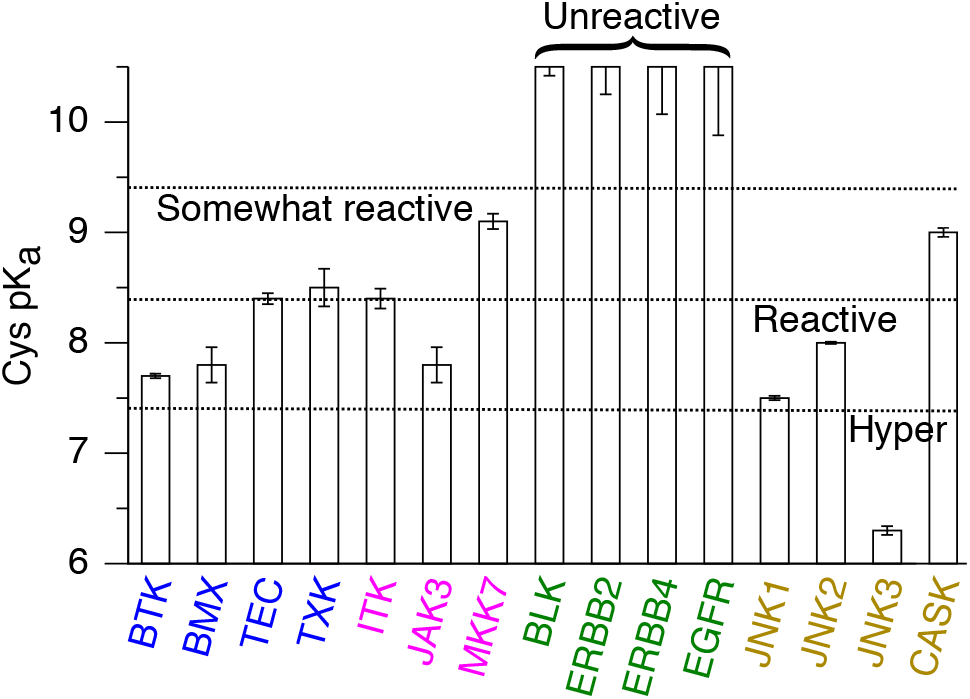
Predicted p*K* _a_’s and reactivities for the FP N-cap and N-cap+2 Cys in 15 kinases. The reactivity definitions are given in Table 2. The exact p*K* _a_ values for the N-cap Cys in EGFR/ERBB2/ERBB4 and BLK are not given, as the thiol form was dominant even at the highest simulation pH 10.5. The kinase names are shown in different colors based on the environment of the FP N-cap Cys: blue (N-cap+3 Asn), magenta (N-cap+3/2 Asp/Glu and N-cap+2/3 Ser/Arg/Lys), and green (N-cap+3 Asp/Glu). The kinases with a FP N-cap+2 Cys are shown in brown.

Sequence alignment of the 11 kinases showed that the FP N-cap Cys is often followed by an Asn or Asp residue at N-cap+3 position (Fig. 1 b and c). A previous experimental study suggested that the N-cap+3 residue may form hydrogen bond (h-bond) or electrostatic interaction with the N-cap residue. ^38^ Our previous computational work ^50,52^ demonstrated that the p*K* _a_ of a Cys can be significantly shifted by h-bonding with proximal residues. Experimental work on various proteins also suggested the role of local h-bonding and electrostatics in stabilizing Cys thiolates. ^39,43,57^ Thus, to understand the varied reactivities of the FP N-cap Cys, we binned the 11 kinases into three groups according to the nearby residues that provide stabilization or destabilization of the N-cap thiolate: Asn at N-cap+3 (BTK/BMX/TEC/TXK); Asp/Glu at N-cap+3/2 and Ser/Arg/Lys at N-cap+2/3 positions (ITK, JAK3, and MKK7); and Asp/Glu at N-cap+3 without other nearby stabilizing h-bond or electrostatic partners (EGFR/ERBB2/ERBB4 and BLK). Below we present analysis of the pH-dependent h-bond and electrostatic interactions to elucidate the FP N-cap Cys reactivities.

### BTK/BMX/TEC/TXK: FP N-cap thiolate stabilized by the N-cap+3 Asn and other h-bond donors

BTK/BMX/TEC/TXK of the Tec family contain an Asn residue at the N-cap+3 position relative to the FP N-cap Cys. The calculated p*K* _a_’s of the N-cap Cys in BTK (C481) and BMX (C496) are 7.7 and 7.8, respectively, indicating that they are nearly hyper-reactive according to our definition (Table 2). Trajectory analysis shows that when deprotonated, Cys481 in BTK accepts h-bonds from both the side chain and backbone amide of Asn484 at N-cap+3 as well as the hydroxyl group and backbone amide of Thr410 (Fig. 3a). The pH-dependent occupancy of the h-bond formation and the pH-dependent deprotonation of Cys481 are perfectly correlated, suggesting that the h-bond interactions contribute to the p*K* _a_ downshift of Cys481 (Fig. 3b), consistent with our previous findings that thiolates tend to be stabilized by nearby h-bond donors. ^50,52^ Similarly, the thiolate form of Cys496 in BMX is stabilized by h-bonding with the sidechain and backbone of Asn499 at N-cap+3 as well as the salt-bridge interaction with Arg540 (Fig. S1).

**Figure 3.**
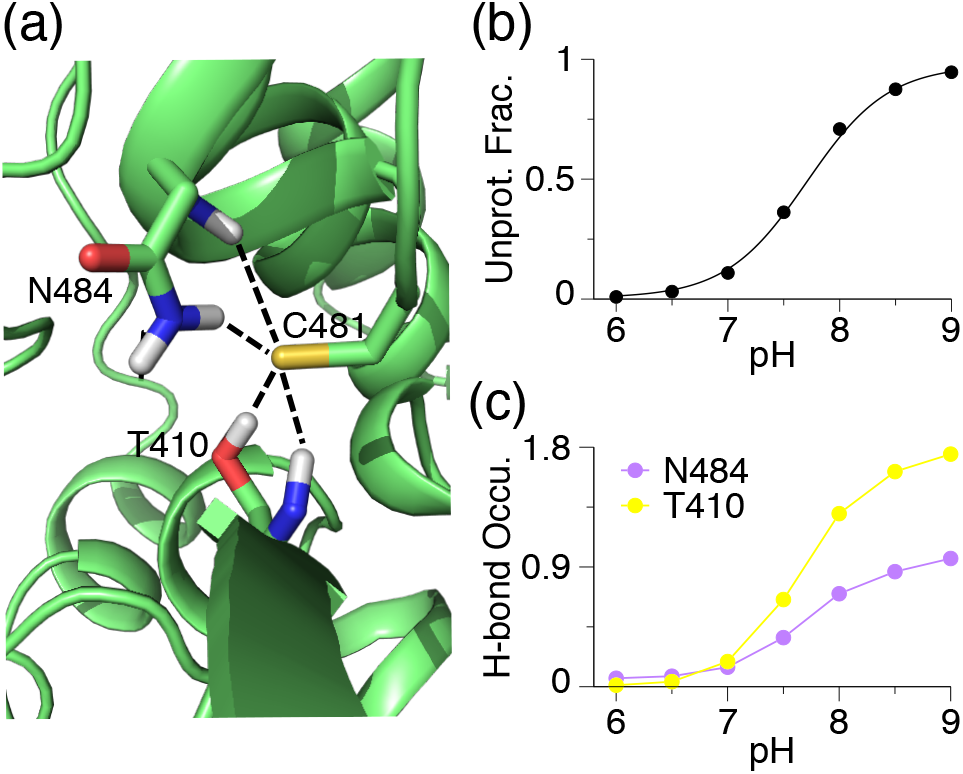
The FP N-cap thiolate is stabilized by hydrogen bonding with N-cap+3 Asn and other residues in BTK. (a) A zoomed-in view of the structural environment of Cys481 in BTK with the Asn484–Cys481 and Thr410–Cys481 h-bonds shown. A snapshot from simulation at pH 9.0 is used. (b) Deprotonated fraction of C481 at different pH. Curve represents the best fit to the Henderson-Hasselbalch equation. (c) Occupancy of the h-bond formation between Cys481 thiolate and Asn484 (sidechain and backbone) or Thr410 (sidechain and backbone) at different pH. A h-bond is defined by the heavy-atom donor-acceptor distance of 3.5 Å and the donor-hydrogen-acceptor angle of 150°.

The calculated p*K* _a_’s of the FP N-cap Cys in TEC (Cys449) and TXK (Cys350) are about 8.5, suggesting that they are somewhat reactive at physiological pH. In TEC, Cys449 thiolate accepts h-bonds from Asn452 at N-cap+3 and Ser378, as well as a salt bridge with Arg493 positioned at N-cap+44. Cys350 in TXK forms h-bonds with Asn353 at N-cap+3 and salt-bridge with Arg394 at N-cap+44. It is interesting to notice that the N-cap+44 Arg which interacts with the N-cap thiolate is the second Arg of the sequence HRDLAARN on the catalytic (C) loop which is highly conserved among protein tyrosine kinases. ^58^ It is also noteworthy that the N-cap Cys does not form h-bond or salt-bridge with i-2 or i-3 residues such as Asn, Ser, Glu, and Arg (Fig. 1c).

### ITK, JAK3, and MKK7: FP N-cap thiolate destabilized by N-cap+3/2 Asp/Glu but stabilized by N-cap+2/3 Ser/Arg/Lys or N-cap+1 Lys/Thr

Several of the N-cap Cys are in close proximity to an anionic side chain (Asp or Glu), which would increase the Cys p*K* _a_ due to electrostatic repulsion. However, h-bonding and salt-bridge formation with the nearby residues can stabilize the thiolate form and thereby decreasing the N-cap Cys p*K* _a_. These opposing effects are experienced by the N-cap Cys in ITK (Cys442), JAK3 (Cys909), and MKK7 (Cys218), resulting in the calculated p*K* _a_’s of 8.4, 7.8, and 9.1, respectively.

Simulations of ITK showed that the deprotonated Cys442 can accept h-bonds from the side chain hydroxyl and backbone amide of Ser444 at N-cap+2 and the backbone amide of Asp445 at N-cap+3, as well as form a salt-bridge with Arg486 positioned at N-cap+44 (Fig. 4a). The excellent correlation between the increasing occupancies of h-bond (or salt bridge) formation and the increasing degree of Cys deprotonation suggests that these interactions stabilize the thiolate form, which compensates for the electrostatic repulsion from the nearby carboxylate group of Asp445 (Fig. 4b and c). As a result, the calculated p*K* _a_ of Cys442 is 8.4, indicating that it is somewhat reactive towards electrophiles.

**Figure 4.**
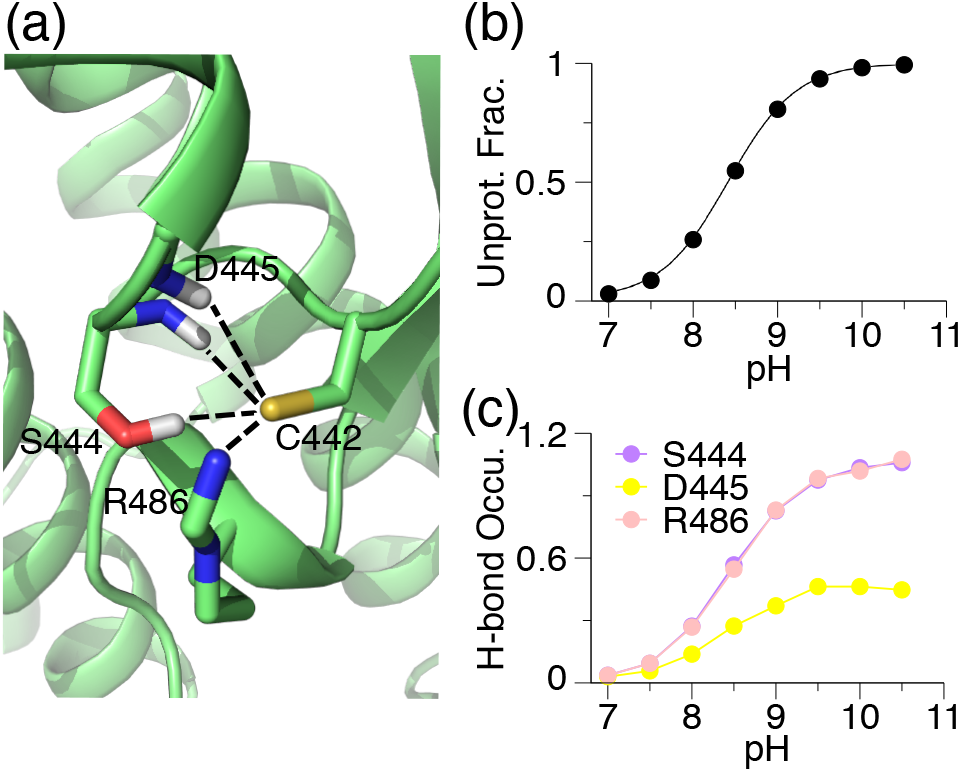
Stabilizing interactions for the FP N-cap Cys thiolate in ITK. (a) Zoomed-in view of the structural environment of Cys442 in ITK taken from the simulation at pH 9.5. (b) pH-dependent deprotonated fraction of the FP Cys442. Curve represents the best fit to the Henderson-Hasselbalch equation. (c) pH-dependent occupancies of the h-bond formation between Cys442 thiolate and the sidechain and backbone of Ser444 (purple), backbone of Asp445 (yellow), and the side chain of Arg486 (light pink).

The compensating interactions are somewhat different for JAK3 and MKK7, in which an attractive electrostatic interaction with a nearby cationic sidechain is also a major contributor. In JAK3, the N-cap Cys909 thiolate is stabilized by salt-bridge interactions with Arg953 at N-cap+44 and h-bonding with Tyr904 at N-cap-5 at pH below pH 8 (Fig. S2). As a result, despite the repulsion with Asp912 at N-cap+3, the p*K* _a_ of Cys909 is downshifted to 7.8, lower than the corresponding N-cap Cys in ITK.

In MKK7, the sidechain-to-sidechain salt-bridge and sidechain-to-backbone h-bond interactions with Lys221 at N-cap+3 as well as the h-bond formation with Ser144 are the stabilizing forces for the thiolate state of the N-cap Cys218 (Fig. S2); these forces compensate for the destabilization due to Glu220 at N-cap+2 position. Overall, the number of stabilizing h-bond/salt bridge interactions with the N-cap Cys in MKK7 is less than those in ITK or JAK3, which may explain why the N-cap Cys p*K* _a_ in MKK7 (9.1) is the highest among the three kinases.

### EGFR/ERBB2/ERBB4 and BLK: FP N-cap thiolate destabilized by N-cap+3 Asp/Glu and desolvation without compensating interactions

The N-cap Cys is buried in EGFR/ERBB2/ERBB4 (of the EGFR family) and BLK (of the Src family) to a similar extent as in the aforementioned kinases; however, there is no nearby h-bond donor or attractive electrostatic interaction partner. Furthermore, the N-cap+3 position is occupied by an anionic Asp (or Glu in ERBB4) which can potentially destabilize the thiolate form of the N-cap Cys. Thus, it is not surprising that the calculated p*K* _a_’s are above 10.5 (the highest simulation pH), suggesting that the N-cap Cys in these kinases are unreactive towards electrophiles.

Curiously, the N-cap Cys in all four kinases have been targeted by the FDA-approved covalent drugs. Most notably, Cys797 in EGFR was the target of the first TCKI, ^5^ and it was subsequently targeted by numerous investigational and FDA-approved inhibitors including ibrutinib, afatinib, and neratinib (Table 1). Analysis of the X-ray structure of EGFR (e.g., PDB ID 4zjv) reveals that the Cys797 thiol may donate a h-bond to the carboxylate group of the N-cap+3 Asp800 (Fig. 5). Similarly, the X-ray structures of ERBB2/ERBB4 and BLK show that the N-cap thiol sulfur is within the h-bond distance to the carboxylate oxygens of N-cap+3 Asp.

**Figure 5.**
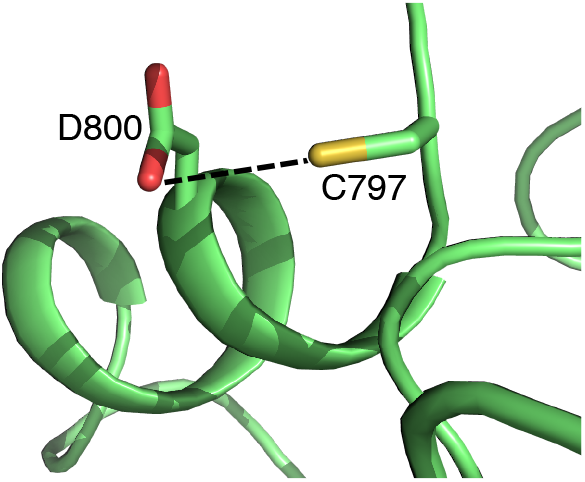
The FP N-cap Cys thiol can donate a h-bond to the N-cap+3 aspartate in EGFR. Zoomed-in view of the structural environment of the N-cap Cys797 in EGFR from the X-ray crystal structure (PDB: 4zjv ^32^). A potential h-bond from Cys797 thiol to the carboxylate of Asp800 is indicated as a dashed line.

The high (calculated) p*K* _a_ of the N-cap and the potential formation of a thiol–aspartate h-bond in the holo structure together suggest that the Michael addition reactions with EGFR/ERBB2/ERBB4 and BLK may proceed via a base-assisted mechanism, in which the N-cap+3 Asp serves as a general base to deprotonate the N-cap Cys in the first stage of the reaction. Indeed, a recent combined quantum mechanical (QM)/MM simulation of EGFR ^59^ demonstrated that in the proximity of a covalent inhibitor carrying an acrylamide warhead, the proton on the N-cap Cys797 thiol can be transferred to the carboxylate of the N-cap+3 Asp, resulting in a stable thiolate/Asp-COOH pair, from which the thiolate then attacks the C*β* of acrylamide generating a carbanion before being reprotonated by the N-cap+3 Asp-COOH to result in the thiol adduct. ^59^ Our result is consistent with this base-assisted mechanism. We should also note that in our apo simulations of the EGFR/ERBB2/ERBB4 and BLK, the thiol group of the N-cap Cys did not form a h-bond with the N-cap+3 Asp/Glu, supporting the hypothesis that the proton transfer only occurs in the presence of an electrophilic warhead. ^59^ Another possibility for the covalent modification of an unreactive Cys is to make use of a base group adjacent to the Michael acceptor warhead. The structure-activity data of EGFR inhibitors suggested that a basic amine, e.g., dialkylamino or pyrrolidine group can abstract a proton from the Cys thiol and thereby catalyzing the reaction. ^30,60,61^

### Other reactive Cys and Lys locations

Since in the CpHMD simulations all Cys and Lys were allowed to titrate simultaneously, we were able to determine their p*K* _a_ values. Table 3 lists the locations of all reactive Cys and Lys predicted by our simulations. Interestingly, a Cys on *β*4 sheet (i-17/18 with respect to the N-cap Cys) exists in 10 kinases and is found reactive or hyper-reactive in BMX/TEC/TXK of the Tec family and ERBB2/EGFR of the EGFR family. The positions of other reactive Cys are proximal to the ATP binding site and have already been targeted by clinical and investigational TCKIs. These locations include the P loop (Cys147 of MKK7), DFG-1 (Cys276 of MKK7), and the A loop (Cys891 of ERBB4).

**Table 3.** Summary of reactive Cys and Lys in the 15 kinases discussed in this work*^a^*

| Kinase | Family | Cys | Location | Reactivity | Major contributors | Lys | Location | Reactivity |
| --- | --- | --- | --- | --- | --- | --- | --- | --- |
| **FP N-cap Cys** |  |  |  |  |  |  |  |  |
| BTK | Tec | **C481** | **front pocket** | reactive | N(+3), T(-71) | K430 | roof | reactive |
| BMX | Tec | C479 | β4 sheet | reactive | | K445 | roof | somewhat |
|  |  | **C496** | **front pocket** | reactive | N(+3), R(+44) |  |  |  |
| | | C549 | β8 sheet | reactive | | | | |
| TEC | Tec | C406 | αC helix | reactive | | | | |
| | | C432 | β4 sheet | reactive | | | | |
|  |  | **C449** | **front pocket** | reactive | S(-71), N(+3), R(+44) |  |  |  |
| TXK | Tec | C333 | β4 sheet | hyper | | | | |
|  |  | **C350** | **front pocket** | somewhat | N(+3), R(+44) |  |  |  |
| | | C402 | β8 sheet | somewhat | | | | |
| ITK | Tec | **C442** | **front pocket** | reactive | S(+2), D(+3), R(+44) |  |  |  |
| JAK3 | JakA | **C909** | **front pocket** | reactive | Y(-5), R(+2), R(+44) | K855 | roof | hyper |
| | | C1028 | αG helix | reactive | | | | |
| MKK7 | STE7 | C147 | **P loop** | hyper |  |  |  |  |
|  |  | **C218** | **front pocket** | reactive | S(-74), K(+3) |  |  |  |
|  |  | C276 | DFG-1 | reactive |  |  |  |  |
| | | C341 | loop (αF/αG) | hyper | | | | |
| EGFR | EGFR | C775 | loop (αC/β4) | reactive | | | | |
| | | C781 | β4 sheet | reactive | | | | |
| ERBB2 | EGFR | C789 | β4 sheet | hyper | | K860 | DFG-3 | reactive |
| ERBB4 | EGFR | C891 | A loop | reactive |  | K858 | DFG-3 | reactive |
| BLK | Src | C460 | loop (αG/αH) | somewhat | | | | |
| **FP N-cap+2 Cys** |  |  |  |  |  |  |  |  |
| JNK1 | MAPK | **C116** | **front pocket** | reactive | Q(+1), S(+39) |  |  |  |
| | | C245 | | reactive | αH helix | | | |
| JNK2 | MAPK | **C116** | **front pocket** | reactive | Q(+1), S(+39) | K153 | β7 sheet | somewhat |
| JNK3 | MAPK | **C154** | **front pocket** | hyper | S(-82), Q(+1), S(+39) |  |  |  |
| | | C283 | | reactive | loop (αG/αH) | | | |
| CASK | CASK | C100 | front pocket | somewhat |  |  |  |  |

<sup>a</sup>The reactivity levels are defined in Table 2. Residues or locations that have been previously targeted by covalent inhibitors are labeled in bold font. Residues that make significant contributions to the $pK_a$ shift of the N-cap Cys are listed with the sequence positions (relative to N-cap) given in the parenthesis. The calculated $pK_a$ 's of all listed residues are given in Table S1.

The local conformational environment of the N-cap Cys is not affected by the conformation of the DFG-motif (in or out) or *α*C helix (in or out). As such, the calculated p*K* _a_ is in-sensitive to the variation of the kinase conformation. For example, using five X-ray structures (in different DFG/*α*C conformations according to KLIFS ^53^), the calculated p*K* _a_ of the N-cap Cys481 is 7.8–8.4 (7.8 with PDB 3pj3, 8.3 with PDB 6o8i, 8.4 with PDB 3piy, 8.3 with PDB 5p9j, and 7.9 with PDB 3oct). In contrast, although the roof catalytic Lys has a p*K* _a_ above 10.4 in the DFG-in or *α*C-in states, the p*K* _a_ value can be significantly downshifted when the kinase adopts the DFG-out/*α*C-out conformation ^50^). Our simulations predicted that the roof Lys in BTK (Lys430, PDB 3pj3), ^25^ BMX (Lys445, PDB 3sxs), ^8^ and JAK3 (Lys855, PDB 5lwm) ^62^ are reactive. Interestingly, although the X-ray structures for these kinases show a DFG out state, *α*C-Glu in BTK and JAK3 is within a salt-bridge distance to the roof Lys. During the simulations, however, the Glu–Lys salt-bridge disrupted, resulting in the sampling of the *α*C-Glu out conformations. The facile transition between the *α*C-Glu in and out positions is consistent with our previous explicit-solvent simulations of the c-Src kinase. ^63^ The Lys at the DFG-3 position has not been targeted before but it is also in the ATP binding site, and the simulations predicted that the DFG-3 Lys in ERBB2 and ERBB4 are reactive, thus providing a new opportunity for covalent targeting. We note that the P-loop and DFG-1 Cys as well as the roof Lys have been previously targeted in other kinases. ^3^

### Comparison to other p*K* _a_ calculations

The CpHMD predicted p*K* _a_’s are in stark contrast to the structure-based calculations based on the empirical PROPKA approach and on the solutions to the PB equation. Most N-cap Cys p*K* _a_’s predicted by the PB calculation (based on the H_++_ server) are between 9.0 and 11.4, similar to those from PROPKA (Table S2). Most notably, while CpHMD simulations gave the p*K* _a_’s of about 7.7–7.8 for the N-cap Cys in BTK, BMX, and JAK3 (the lowest among the 11 kinases, Table S1), the PROPKA calculated p*K* _a_’s are 9.6, 9.9, 10.3, and 9.0, and those from PB are 9.6, 9.9, 8.8, and 9.0, respectively (Table S2). The large discrepancy, which has been noted in our previous benchmark studies, ^48,50^ can be attributed to the h-bonds and electrostatic interactions that are absent in the X-ray structures but emerge from the pH-dependent conformational dynamics captured by CpHMD and not by structure-based calculations. ^64^ For example, in BTK, the sidechain hydroxyl and backbone amide of Thr410 donate h-bonds to stabilize the N-cap Cys481 in a pH-dependent manner (Fig. 3); however, these interactions are absent in the X-ray structure and therefore not accounted for by the PROPKA or PB calculations. Similarly, in ITK, the stabilizing interactions with Ser444 and Arg486 (Fig. 4) are absent in the X-ray structure and therefore the p*K* _a_’s from the PROKA and PB calculations are over 10.4, two units higher than the value of 8.4 from the CpHMD titration.

The TI simulations and alternative constant pH methods based on the hybrid-solvent Monte-Carlo (MC)/MD and non-equilibrium MD/MC schemes of Rowley and coworker ^65^ gave the p*K* _a_’s of 10.4–13.0 for the N-cap Cys in BTK/BMX/ITK, JAK3, and 11.1–13.5 for EGFR (Table S2). Except for EGFR, these results are in disagreement with our CpHMD simulations which suggested that all four N-cap Cys are reactive with the p*K* _a_’s of 7.7–8.4. With regards to the N-cap Cys797 in EGFR, Rowley and coworker calculated its p*K* _a_ with either protonated or deprotonated Asp800. Interestingly, the calculated p*K* _a_ was above 10 in both cases. ^65^

### The FP N-cap+2 Cys in JNKs and CASK

JNK1/JNK2/JNK3 of the MAPK family and CASK of the CASK family possess a Cys similar in position as the N-cap Cys discussed so far. This N-cap+2 Cys, Cys116 in JNK1/JNK2, Cys154 in JNK3, or Cys100 in CASK is the second N-terminal residue in the *α*D helix, and has an Ile in the N-cap+3 position instead of Asn/Asp/Glu. The calculated p*K* _a_’s for Cys116 in JNK1/JNK2 and Cys154 in JNK3 are 7.5, 8.0, and 6.3 respectively, while that for Cys100 in CASK is 9.0 (Table S2). Analysis showed that in JNK1 and JNK2, Cys116 thiolate can accept h-bond formation with Gln117 and Ser155. In JNK3, Cys154 thiolate is additionally stabilized by the h-bond formation with Ser72, which may explain the lower p*K* _a_ (hyper-reactivity) of Cys154 compared to Cys116 in JNK1/JNK2. As to CASK, the deprotonation of Cys100 is mainly correlated with the salt-bridge formation with Arg302 and to a lesser extent with the h-bond formation with Asn299. The number of h-bonds formed by Cys100 in CASK is lower than that by Cys116 in JNK1/JNK2 or Cys154 in JNK3. As a result, the p*K* _a_ upshift due to desolvation is not offset, leading to a higher p*K* _a_ value of Cys100 as compared to Cys116 or Cys154 in JNKs.

## CONCLUDING REMARKS

N-cap Cys has long been hypothesized as hyper-reactive due to the thiolate-helix dipole interactions. ^38,39^ In support of this hypothesis, the N-cap Cys in all human kinases have been covalently targeted. Our data demonstrated that the N-cap position alone may not stabilize the thiolate form making the Cys reactive at physiological pH. CpHMD simulations confirmed the reactivities of the N-cap Cys in BTK/BMX/TEC/TXK/ITK and JAK3, with the calculated p*K* _a_’s ranging from 7.7 to 8.5. The calculated p*K* _a_’s of 7.7 and 8.4 for BTK and ITK are in excellent agreement with the respective experimental values of 7.7 and 8.5. ^55^ In comparison, the calculated p*K* _a_ is 9.1 for the N-cap Cys in MKK7, suggesting a lower reactivity compared to the aforementioned Tec family kinases and JAK3.

Surprisingly, the calculated p*K* _a_’s are above 10.4 for the N-cap Cys in EGFR/ERBB2/ERBB4 and BLK, suggesting that the thiol form is predominant at physiological pH despite the N-cap position. Curiously, the calculated p*K* _a_ for EGFR is 5 units higher than that from the pH-dependent bromobimane-thiol reaction rate. This discrepancy cannot be explained by the lack of polarization of additive force field based CpHMD simulations, which may lead to a small underestimation of the helix dipole ^66^ and consequently the p*K* _a_ downshift due to the thiolate-dipole interaction. We suggest that the thiol addition reaction of EGFR and ERBB2/ERBB4 as well as BLK may be initiated from a thiol state by following a base-assisted mechanism. EGFR/ERBB2/ERBB4 and BLK all contain an Asp in the N-cap+3 position relative to the N-cap Cys, and in the holo crystal structures, this Asp is within h-bond distance to the Cys. A recent QM/MM study suggested that in the proximity of an inhibitor, a Cys Asp h-bonded pair may form, from which the thiolate then attacks the acrylamide warhead on the inhibitor. ^59^ Thus, it is possible that the Cys797 p*K* _a_ determined from the bromobimane-thiol reaction. ^56^ refers to the Cys797 Asp800 pair in the presence of bromobimane.

Other base-assisted mechanisms are also possible for thiol-Michael addition reactions. Several experimental studies of the structure-activity relationships of EGFR inhibitors suggested that an amine group proximal to the electrophilic warhead may act as a general base to abstract the proton from the N-cap thiol yielding the nucleophilic thiolate. ^30,60,61^ Additionally, a nearby water may in principle serve as a general base. Finally, in the absence of a general base, a direct addition mechanism is conceivable, whereby in the first step the carbonyl oxygen abstracts the proton from the N-cap thiol, as proposed by a most recent QM/MM study of BTK modification by ibrutinib assuming the N-cap Cys is protonated. ^67^

A corollary to the N-cap thiolate hypothesis is that a Cys placed further down the helical sequence, e.g., at N2 or N3 position, should have a higher p*K* _a_. The CpHMD simulations of JNK1/JNK2/JNK3 and CASK showed that this needs not be the case. The calculated p*K* _a_’s of the N-cap+2 Cys in JNKs range from 6.3 to 8.0, consistent with the fact that they have all been covalently modified. ^17,18^ The calculated p*K* _a_ of N-cap+2 Cys in CASK indicates it is somewhat reactive; no covalent inhibitor targeting this position has been reported yet.

Our analysis showed that while the solvent accessibility of the N-cap or N-cap+2 Cys in various kinases is very similar (all largely buried), the h-bond and electrostatic environment can be very different, which gives rise to the varied Cys p*K* _a_’s. Specifically, interaction with nearby h-bond donors is a major determinant for the p*K* _a_ downshift, while solvent exclusion is a major determinant of the p*K* _a_ upshift, consistent with our previous findings in the context of other proteins. ^50,52^ Notably, the presence of a h-bond donor at the N-cap+3 position (e.g., Asn) to the stabilizes the thiolate form, e.g., in BTK/BMX/TXK. By contrast, the presence of an anionic sidechain (e.g. Asp or Glu) at the N-cap+3 position destabilizes the thiolate form. Although ITK, JAK3, BLK, and EGFR/ERBB2/ERBB4 all have an N-cap+3 Asp next to the N-cap Cys, only ITK and JAK3 have favorable h-bond interactions that stabilize the thiolate form. As a result, the N-cap Cys in ITK and JAK3 are reactive, whereas those in BLK and EGFR/ERBB2/ERBB4 are unreactive.

A unique capability of CpHMD simulations is to capture pH-dependent formation of h-bond and electrostatic interactions that are nascent or absent in the static (X-ray) structure but can nonetheless significantly impact the protonation state. Such interactions with the N-cap or N-cap+2 Cys cannot be accounted for by traditional structure-based PB or empirical calculations; the predicted high p*K* _a_’s are dominated by the large contribution from the desolvation penalty.

The main caveats of the current CpHMD approach include the neglect of polarization, which may lead to a small under-estimation of the Cys p*K* _a_ downshift due to underestimation of the helix dipole, as well as the neglect of proton transfer between Cys and a nearby general base. QM studies will be conducted in the future to further test the hypothesis that the base-assisted mechanisms underlie the Michael addition reactions with the N-cap Cys in EGFR/ERBB2/ERBB4 and BLK. Taken together, our work offers a systematic understanding of cysteine reactivities in kinases and demonstrates the utility of GPU-accelerated CpHMD simulations in making accurate predictions and rationalization of Cys reactivities to guide the design of targeted covalent inhibitors.

## METHODS AND PROTOCOLS

### Structure Preparation

Except for TEC, TXK, and BLK, the initial structures were taken from the Protein Data Bank (PDB): BTK (3pj3), ^25^ BMX (3sxs), ^8^ ITK (3miy), ^24^ JAK3 (5lwm), ^62^ EGFR (4zjv), ^32^ ERBB2 (3pp0), ^31^ ERBB4 (2r4b), ^30^ JNK1 (2xrw), ^68^ JNK2 (3e7o), ^69^ JNK3 (6emh), ^70^ and CASK (3mfr). ^71^ The structures of TEC, TXK, and BLK were built using SWISS-MODEL. ^20^ For each structure, the open source tool pdbfixer from the OpenMM package ^72^ was used to remove non-protein heavy atoms and to add missing residues, atoms, acetylated N terminus and amidated C terminus, disulfide bonds (if present), and hydrogen atoms. Lys and Cys had one dummy hydrogen, while His had two dummy hydrogens.

### Simulation Protocol

The generalized Born based CpHMD simulations ^47,48^ were performed using the pmemd engine of AMBER18 (AMBER 2018). ^46^ The pH replica-exchange titration simulations were performed on the GPUs using our asynchronous replica-exchange implementation. ^51^ The proteins were represented by the ff14sb protein force field, ^73^ and solvent represented by the GBNeck2 (igb=8) implicit solvent model ^49^ with mbondi3 intrinsic Born radii and 0.15 M ionic strength. Modifications to the intrinsic Born radii were made for His and Cys sidechains, as discussed in our previous work. ^47,48,50^ Prior to the CpHMD runs, energy minimization was performed in the GB solvent using 500 steps of steepest descent and 500 steps of the conjugate gradient method with a harmonic force constant of 50 kcal/mol/Å^2^ applied to the heavy atoms. Following minimization, four stages of equilibration were performed at 300 K using 2000 steps for each stage. A harmonic force constant applied to the heavy atoms was gradually decreased from 5.0, 2.0, 1.0, to 0 kcal/mol/Å^2^.

One set of pH replica-exchange CpHMD simulations was carried out for each protein. For BTK, ITK, and ERBB4, 8 independent pH titrations were also performed at pH conditions from 6.0 to 9.5, with a 0.5 unit interval. Each pH condition was run for 10 ns. These simulations gave very similar results as the replica-exchange runs. The pH conditions in the replica-exchange simulations were from 7.0 to 10.5, except BTK which had a pH range of 6.0–9.0 and MKK7 which had a pH range of 7.0–11.0. The pH conditions were placed at every 0.5 unit, except for MKK7 which had an extra pH condition of 9.25. Bonds involving hydrogen atoms were constrained with the SHAKE algorithm allowing a 2-fs timestep. Exchanges between adjacent pH replicas were attempted every 1000 steps (2 ps) according to the Metropolis criterion. ^74^ Each replica lasted 50 ns (aggregate time of 400 ns) for BTK, BMX, TXK, ITK, JAK3, MKK7, and 30 ns (aggregate time of 240 ns) for the other kinases (these proteins had faster convergence). Each replica underwent Langevin dynamics at 300 K with an effectively infinite cutoff (999 Å) for nonbonded interactions. All side chains of His, Cys, and Lys were allowed to titrate, with the experimental model alanine penta-peptide p*K* _a_’s of 6.5, 8.5, and 10.4, respectively. ^45^ If not stated, all settings were identical to our previous work. ^47,50,52^

### p*K* _a_ calculations

The unprotonated fraction (*S*) of a titratable residue at different pH values was calculated using the definitions of the protonated (*λ* < 0.2) and deprotonated (*λ* > 0.8) states, as in our previous work. ^47,48,52^ The p*K* _a_ was calculated by fitting the *S* values to the generalized Henderson Hasselbalch (HH) equation, *S* = 1/(1 + 10^*n*(*pK*_a_−pH)^), where *n* is the Hill coefficient.

The bootstrap method was used to estimate the errors of the calculated p*K* _a_’s. ^48^ The *S* values based on the 10-ns simulations at different pH were combined to generate m^*n*^ sets of titration data, where *m* is the number of the 10 ns simulations per pH and *n* is the number of pH conditions. For each set of titration data (*S* vs pH), a p*K* _a_ value was calculated. An average and standard deviation of all the calculated p*K* _a_’s are reported (Fig. 2 and Table S1).

## Supporting Information Available

Supplemental tables and figures are included.

## ACKNOWLEDGEMENT

The authors acknowledge National Institutes of Health (R01GM098818 and R01CA256557) for funding.

