## Supplemental tables and figures for "Reactivities of the Front Pocket N-Cap Cysteines in Human Kinases"

#### List of Tables

S1 Summary of the CpHMD calculated p$K_a$'s of all reactive Cys and Lys in the 15 kinases studied in this work . . . . . . . . . . . . . . . . . . . . . . . . . . S-3

S2 Front-pocket N-cap and N-cap+2 Cys p$K_a$'s calculated with other popular computational methods . . . . . . . . . . . . . . . . . . . . . . . . . . . . S-4

#### List of Figures


S1 Titration curves of the FP N-cap Cys and pH-dependent hydrogen bond and salt-bridge interactions in BTK, BMX, and TEC . . . . . . . . . . . . . . . S-5

S2 Titration curves of the FP N-cap Cys and pH-dependent hydrogen bond and salt-bridge interactions in ITK, JAK3 and MKK7. . . . . . . . . . . . . . . S-6

S3 Titration curves of the FP N-cap Cys and pH-dependent hydrogen bond and salt-bridge interactions in JNK1, JNK2, JNK3 and CASK. . . . . . . . . . S-7

S4 Titration and h-bond formation of FP Cys in BLK, ERBB2, ERBB4, and EGFR. . . . . . . . . . . . . . . . . . . . . . . . . . . . . . . . . . . . . . . . . S-8

S5 Titration of other reactive Cys in BMX, TEC, TXK, JAK3,JNK1, JNK3, MKK7, BLK, ERBB2, EGFR, ERBB4. . . . . . . . . . . . . . . . . . . . . . . . . . . . S-9

S6 Titration of reactive Lys in BTK, BMX, JAK3, ERBB2, ERBB4, and JNK2. . S-10

#### Supplemental tables

Table S1: Summary of the CpHMD calculated p$K_a$'s of all reactive Cys and Lys in the 15 kinases studied in this work

| Kinase | Cys[a] | Location | p$K_a$[b] | boot-strap error | Lys | Location | p$K_a$ |
|---|---|---|---|---|---|---|---|
| BTK | **C481** | **FP N-cap** | 7.7 | 0.02 | K430 | roof | 7.5 |
| BMX | C479 | $\beta$4 | 7.5 | | K445 | roof | 8.7 |
| | **C496** | **FP N-cap** | 7.8 | 0.16 | | | |
| | C549 | $\beta$8 | 7.8 | | | | |
| TEC | C406 | $\alpha$C-helix | 8.4 | | | | |
| | C432 | $\beta$4 | 7.6 | | | | |
| | **C449** | **FP N-cap** | 8.4 | 0.05 | | | |
| TXK | C333 | $\beta$4 | 6.5 | | | | |
| | **C350** | **FP N-cap** | 8.5 | 0.17 | | | |
| | C402 | $\beta$8 | 9.3 | | | | |
| ITK | **C442** | **FP N-cap** | 8.4 | 0.09 | | | |
| JAK3 | **C909** | **FP N-cap** | 7.8 | 0.16 | | | |
| | C1028 | $\alpha$G helix | 7.7 | | K855 | roof | 7.3 |
| MKK7 | C147 | P loop | 7.1 | | | | |
| | **C218** | **FP N-cap** | 9.1 | 0.07 | | | |
| | C276 | head of DFG-1 | 8.2 | | | | |
| | C341 | $\alpha$F/$\alpha$G helix | 7.2 | | | | |
| EGFR | C775 | $\alpha$C-helix/$\beta$4 | 7.9 | | | | |
| | C781 | $\beta$4 sheet | 7.7 | | | | |
| EBRR2 | C789 | $\beta$4 sheet | 7.4 | | K860 | DFG-3 | 8.2 |
| EBRR4 | C891 | A loop | 8.2 | | K858 | DFG-3 | 8.3 |
| BLK | C460 | $\alpha$G/$\alpha$H helix | 9.0 | | | | |
| JNK1 | **C116** | **FP N-cap+2** | 7.5 | | | | |
| | C245 | $\alpha$G/$\alpha$H helix | 7.8 | | | | |
| JNK2 | **C116** | **FP N-cap+2** | 8.0 | | K153 | $\beta$7 sheet | 8.9 |
| JNK3 | **C154** | **FP N-cap+2** | 6.3 | | | | |
| | C283 | $\alpha$G/$\alpha$H helix | 7.6 | | | | |
| CASK | **C100** | **FP N-cap+2** | 9.0 | | | | |

[a] Front pocket (FP) N-cap and N-cap+2 cysteines are labeled in bold font. [b] The converged part of the simulation was used for p$K_a$ calculations and analysis. The data from 30 ns–50 per replica was used for BMX due to slower convergence. For other kinases, data from 10–30 ns or 20–30 ns per replica was used. Boot-strap errors are listed for the FP N-cap Cys.

Table S2: Front-pocket N-cap and N-cap+2 Cys p$K_\mathrm{a}$'s calculated with other popular computational methods

| Kinase | Family | Group | PDB | Residue | CpHMD | PROPKA$^a$ | H++$^b$ | RETI$^c$ | pH-REMD$^d$ | neMD/MC$^e$ |
|---|---|---|---|---|---|---|---|---|---|---|
| BTK | Tec | TK | 3pj3 | C481 | 7.7 | 9.6 | 9.6 | 10.41±0.80 | | |
| BMX | Tec | TK | 3sxs | C496 | 7.8 | 9.9 | 9.9 | 10.31±0.46 | | |
| TEC | Tec | TK | 4hct | C449 | 8.4 | 9.2 | 8.3 | | | |
| ITK | Tec | TK | 3miy | C442 | 8.4 | 10.7 | 10.4 | 11.96±0.55 | | |
| TXK | Tec | TK | 3t9t | C350 | 8.5 | 10.4 | 9.7 | | | |
| JAK3 | JakA | TK | 5lwm | C909 | 7.8 | 10.3 | 8.8 | 13.0±0.4 | 12.7 | 11.1 |
| EGFR | EGFR | TK | 4zjv | C797 | 10.5 | 11.4 | 10.8 | 11.1±0.7 | 13.5 | 11.5 |
| ERBB2 | EGFR | TK | 3pp0 | C805 | 10.5 | 10.8 | 10.9 | | | |
| ERBB4 | EGFR | TK | 2r4b | C803 | 10.5 | 11.0 | 9.8 | | | |
| BLK | Src | TK | 4mxy | C319 | 10.5 | 10.6 | 11.4 | | | |
| MKK7 | STE7 | STE | 6qfr | C218 | 9.1 | 9.1 | 9.7 | | | |
| JNK1 | MAPK | CMGC | 2xrw | C116 | 7.5 | 10.0 | 9.6 | | | |
| JNK2 | MAPK | CMGC | 3e7o | C116 | 8.0 | 10.2 | 9.7 | 7.0±0.8 | | |
| JNK3 | MAPK | CMGC | 6emh | C154 | 6.3 | 10.4 | 10.2 | | | |
| CASK | CASK | CAMK | 3mfr | C100 | 9.0 | 13.0 | >12.0 | | | |

$^a$ Calculations performed with PROPKA3.[S1] $^b$ Calculations performed with H++3.0 web server.[S2] $^{c,d,e}$ Calculations using the replica-exchange thermodynamics integration (RETI) simulations and the two Monte-Carlo (MC)/molecular dynamics (MD) methods from the work of Awoonor-Williams and Rowley.[S3] pH-REMD refers to the replica-exchange constant pH MD method that employs MC protonation steps in implicit solvent and replica-exchange MD sampling in explicit solvent.[S4] neMD/MC refers to the Gibbs sampling method that combines non-equilibrium MD simulations in explicit solvent with MC moves in explicit solvent.[S5]

### Supplemental figures

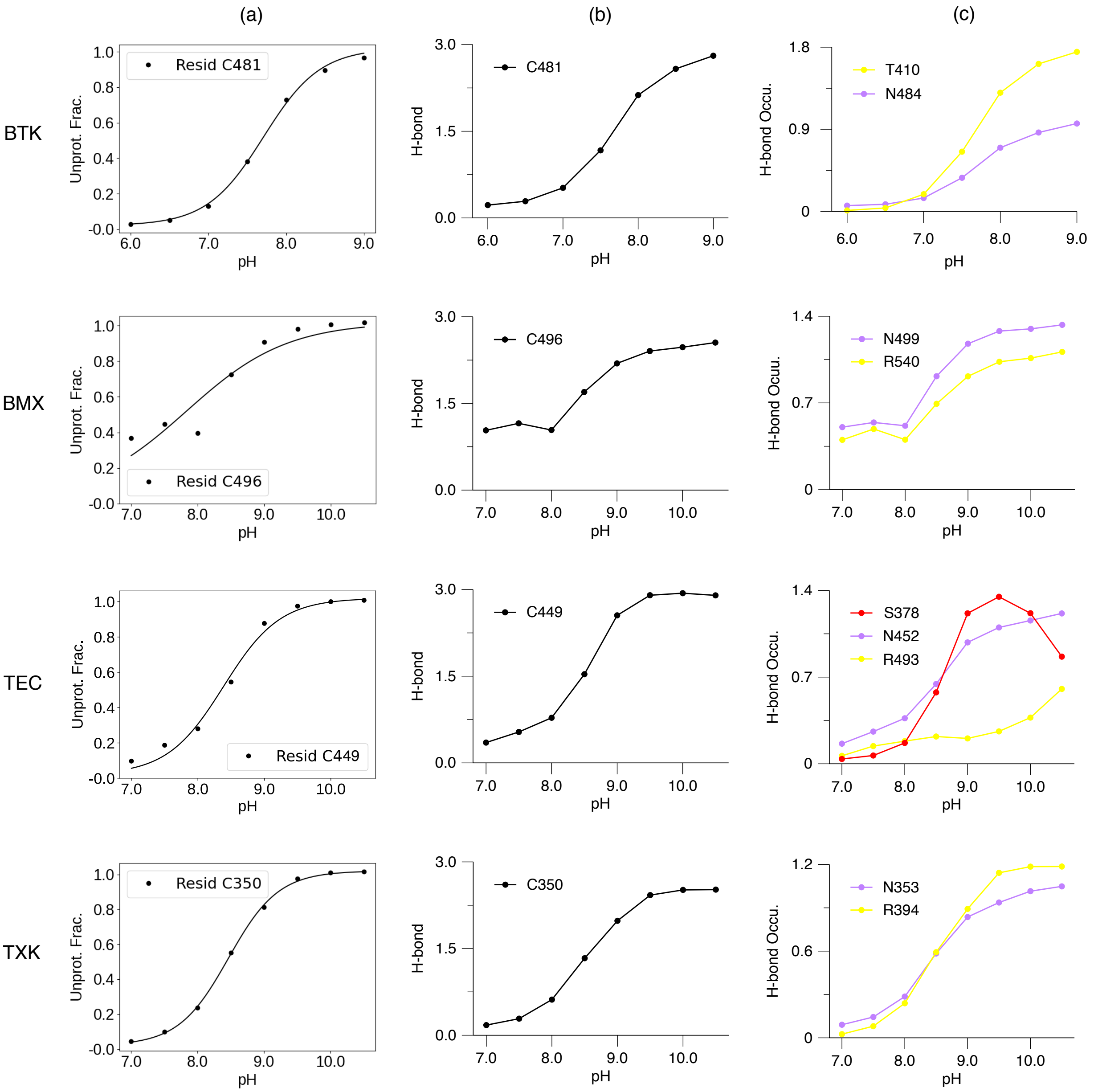


Figure S1: **Titration of the FP N-cap Cys and local hydrogen bonding and electrostatic interactions in BTK, BMX, and TEC.** (a) pH-dependent deprotonated fractions of C481 in BTK, C496 in BMX, C449 in TEC, and C350 in TXK. Curve represents the best fit to the Henderson-Hasselbalch equation. (b) pH-dependent hydrogen bond formation. (c) pH-dependent occupancy of the hydrogen bond and salt bridge formation. The latter used the same distance cutoff. A hydrogen bond is considered present if the heavy-atom donor-acceptor distance is below 3.5 Å and the donor-hydrogen-acceptor angle greater than 150°.

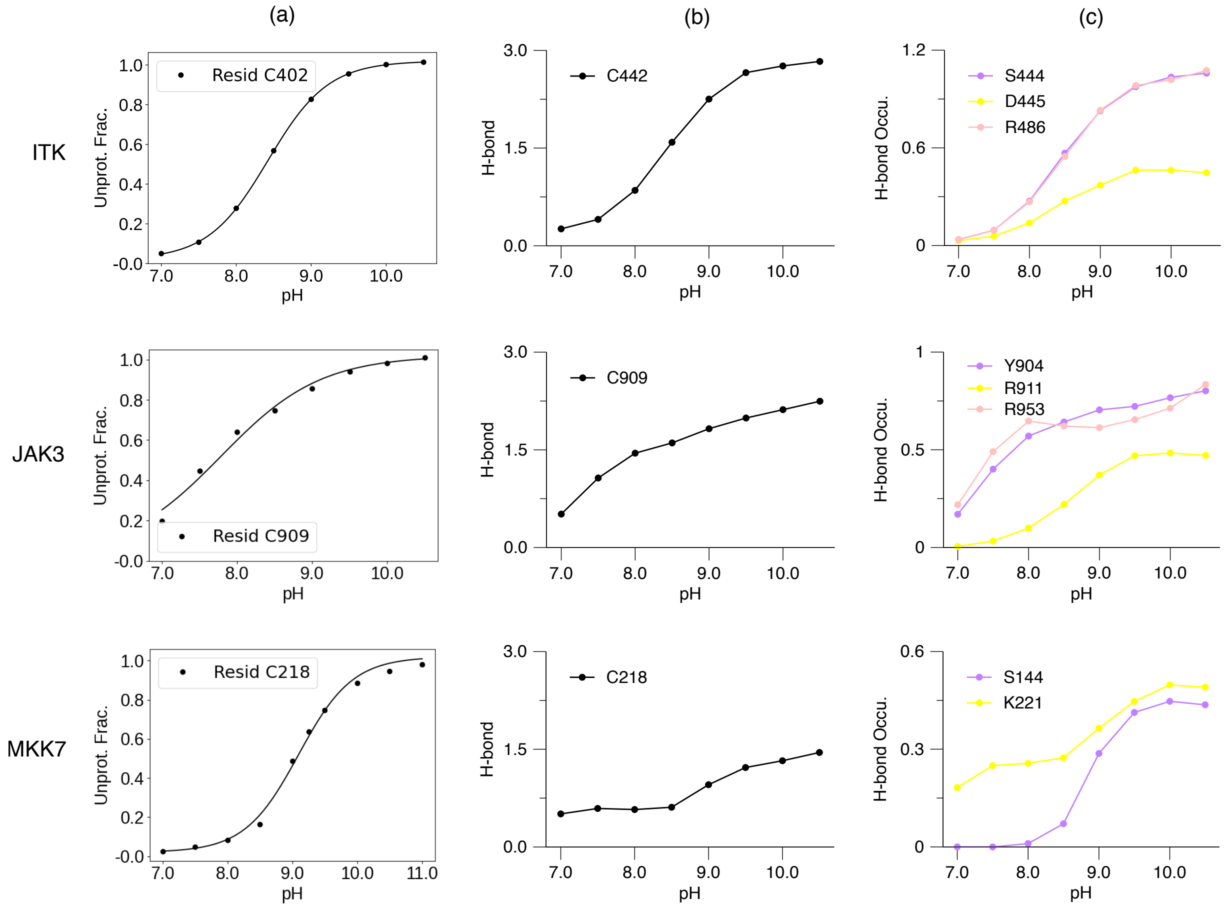


Figure S2: **Titration of the FP N-cap Cys and local hydrogen bonding and electrostatic interactions in ITK, JAK3 and MKK7.** (a) pH-dependent deprotonated fractions of C442 in ITK, C909 in JAK3, and C218 in MKK7. Curve represents the best fit to the Henderson-Hasselbalch equation. (b) pH-dependent h-bond formation of the front-pocket N-cap Cys. (c) pH-dependent occupancy of the hydrogen bonds. A hydrogen bond is considered present if the heavy-atom donor-acceptor distance is below 3.5 Å and the donor-hydrogen-acceptor angle greater than 150 degree.

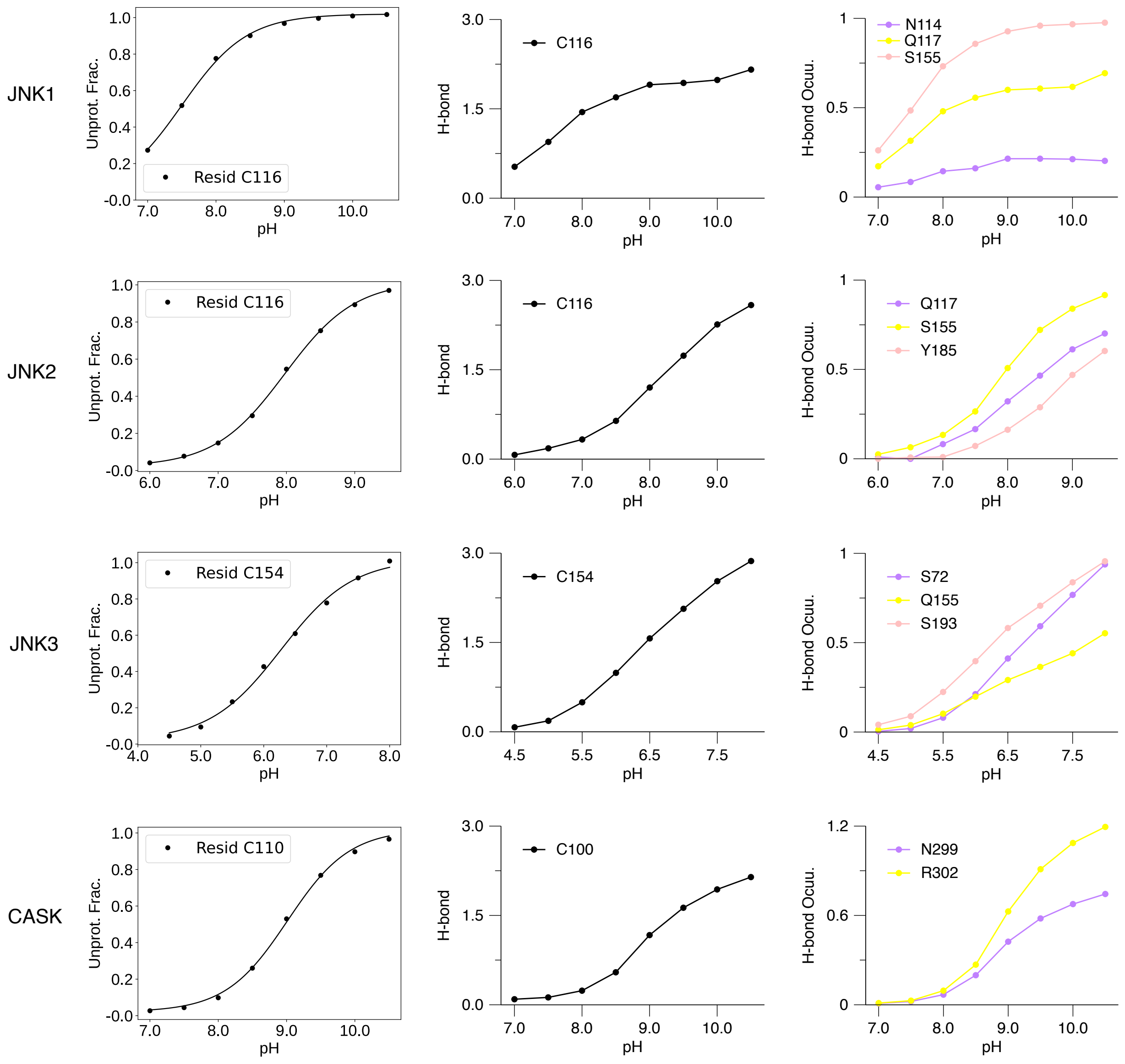


Figure S3: **Titration of the FP N-cap Cys and local hydrogen bonding and electrostatic interactions in JNK1, JNK2, JNK3 and CASK.** (a) pH-dependent deprotonated fractions of C116 in JNK1, C116 in JNK2, C154 in JNK3, and C110 in CASK. Curve represents the best fit to the Henderson-Hasselbalch equation. (b) pH-dependent h-bond formation of the front-pocket N-cap Cys. (c) pH-dependent occupancy of the hydrogen bonds. A hydrogen bond is considered present if the heavy-atom donor-acceptor distance is below 3.5 Å and the donor-hydrogen-acceptor angle greater than 150 degree.

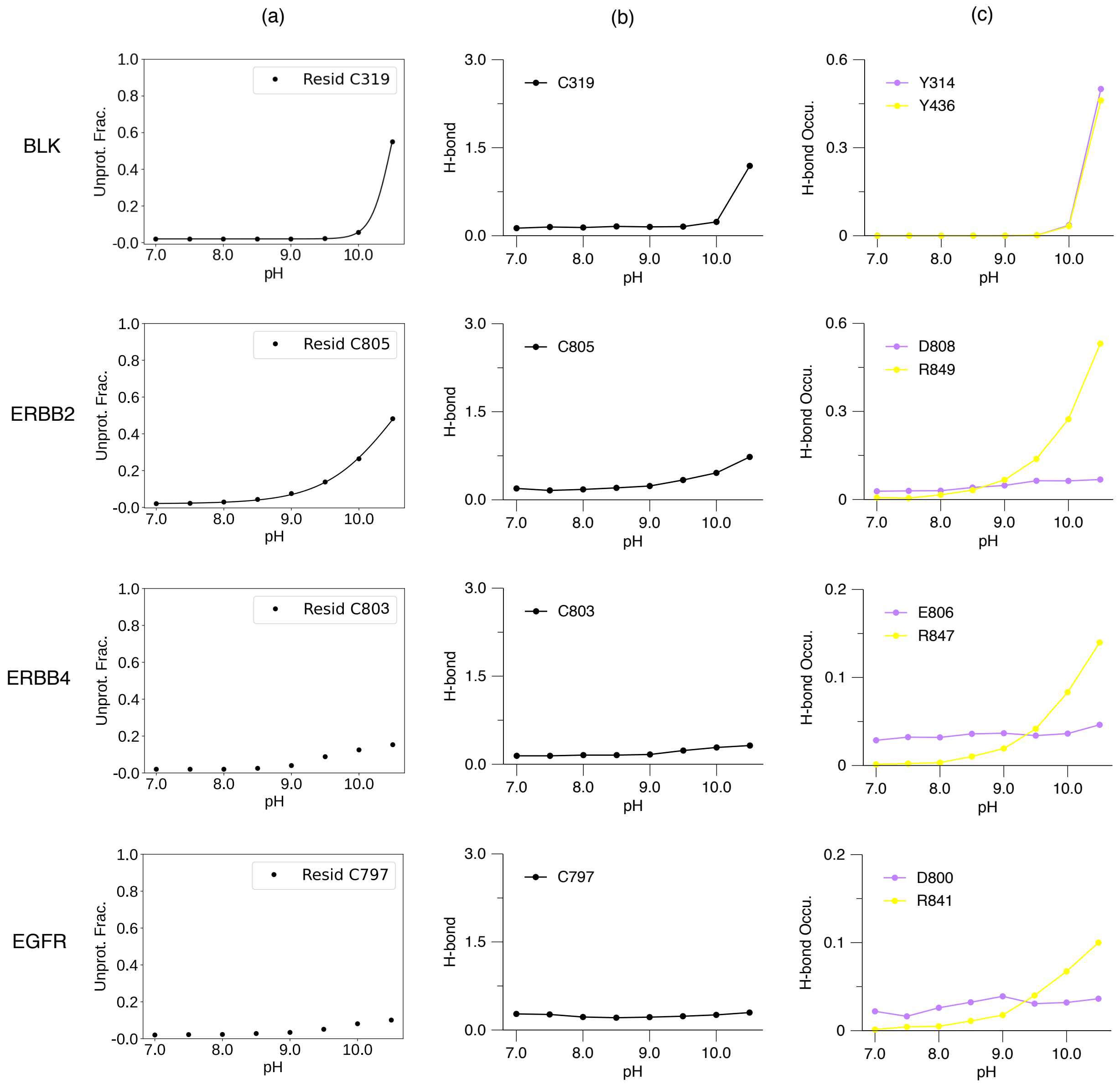


Figure S4: **Titration and h-bond formation of FP Cys in BLK, ERBB2, ERBB4, and EGFR.** (a) pH-dependent deprotonated fractions of C319 in BLK, C805 of ERBB2, C803 of ERBB4 and C797 in EGFR. Curve represents the best fit to the Henderson-Hasselbalch equation. (b) pH-dependent h-bond formation of the front-pocket Cys. (c) pH-dependent occupancy of the hydrogen bonds. A hydrogen bond is considered present if the heavy-atom donor-acceptor distance is below 3.5 Å and the donor-hydrogen-acceptor angle greater than 150°.

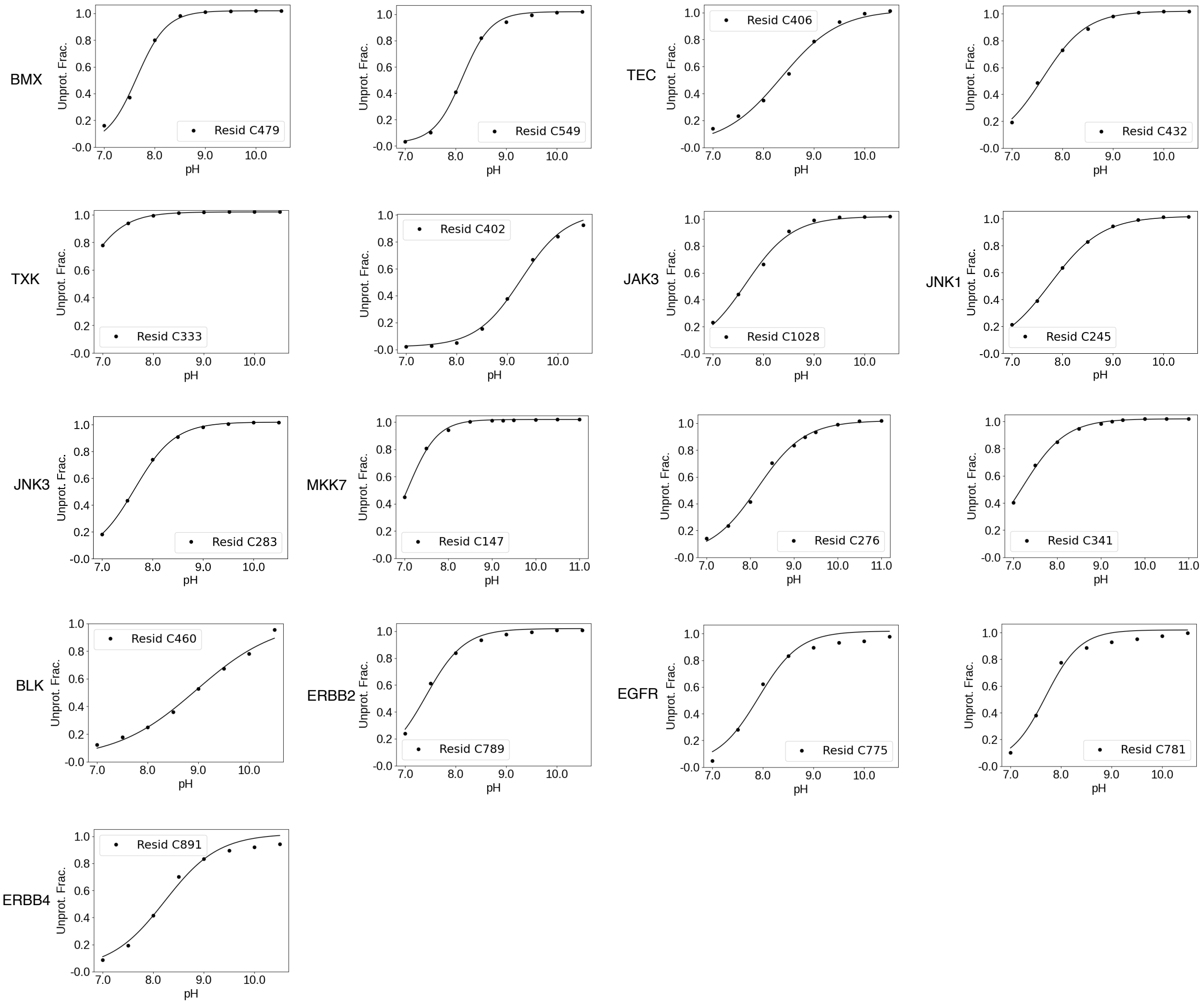


Figure S5: **Titration of other reactive Cys in BMX, TEC, TXK, JAK3, JNK1, JNK3, MKK7, BLK, ERBB2, EGFR, ERBB4.** pH-dependent deprotonated fractions of C479 and C549 in BMX, C406 and C432 in TEC, C333 and C402 in TXK, C1028 in JAK3, C245 in JNK1, C283 in JNK3, C147, C276 and C341 in MKK7, C460 in BLK, C789 in ERBB2, C775 and C781 in EGFR, C891 in ERBB4. Curve represents the best fit to the Henderson-Hasselbalch equation.

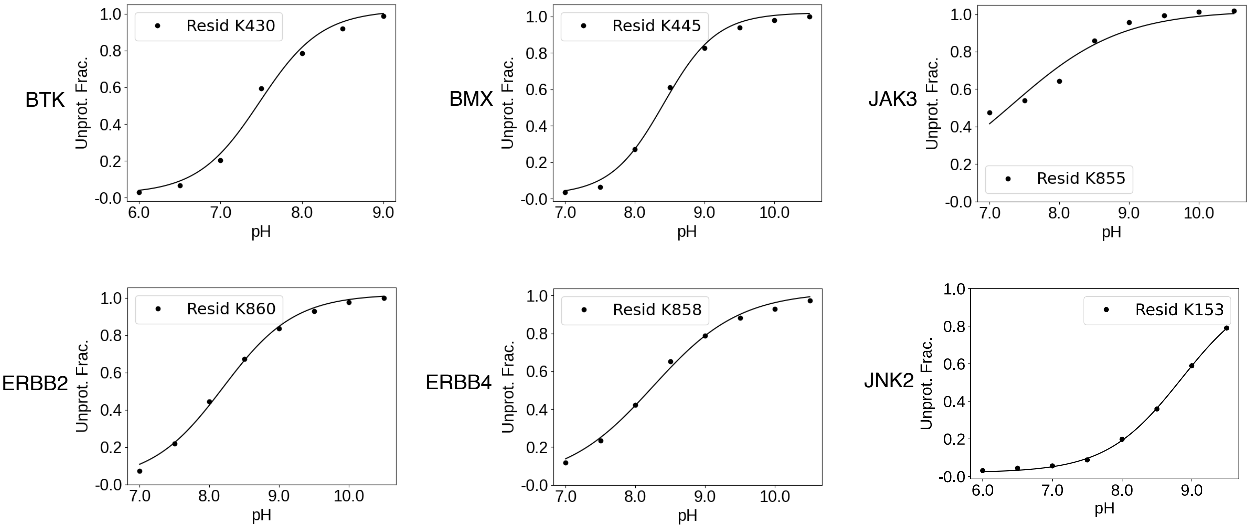


Figure S6: **Titration of reactive Lys in BTK, BMX, JAK3, ERBB2, ERBB4, and JNK2** pH-dependent deprotonated fractions of K430 in BTK, K445 in BMX, K855 in JAK3, K860 in ERBB2, K858 in ERBB4, and K153 in JNK2. Curve represents the best fit to the Henderson-Hasselbalch equation.
